# Enteropathogenic *Escherichia coli* manipulates the host exocyst complex to enhance pedestal formation

**DOI:** 10.1101/2025.09.18.677024

**Authors:** Thilina U.B. Herath, Pasan Dahanayake, Roman Mortuza, Antonella Gianfelice, Attika Rehman, Mihnea Bostina, Wanyin Deng, Andrew S. Santos, B. Brett Finlay, Keith Ireton

**Affiliations:** Department of Microbiology and Immunology, University of Otago, Dunedin, New Zealand; Michael Smith Laboratories and Department of Microbiology and Immunology, University of British Columbia, Vancouver, Canada; Department of Microbiology and Immunology, University of British Columbia, Vancouver, Canada

## Abstract

Enteropathogenic *Escherichia coli* (EPEC), a major cause of diarrheal disease, produces plasma membrane pedestals that promote colonization of host cells. Critical for pedestal formation is EPEC’s type III secretion system, which injects ∼ 20 effector proteins into human cells. One of these effectors is Tir, which inserts into the host plasma membrane and stimulates the assembly of actin filaments essential for pedestal generation. To date, actin polymerization is the only host process known to contribute to pedestal formation. Here we report that EPEC co-opts the membrane trafficking pathway of polarized exocytosis, which acts together with actin polymerization to allow the efficient production of pedestals. Polarized exocytosis is mediated by the exocyst — a human octameric complex that uses intracellular vesicles to expand the plasma membrane. We found that EPEC stimulated exocytosis at sites of pedestal formation in a manner dependent on the exocyst. The bacterial effector EspH recruited the exocyst and promoted exocytosis. Genetic inactivation of *espH* or RNA interference (RNAi)-induced depletion of exocyst components reduced both the frequency and size of pedestals. Additional RNAi experiments indicated that the exocyst is dispensable for actin filament assembly in pedestals. Co-depletion of components of the exocyst and the Arp2/3 complex showed that exocytosis and actin polymerization make additive contributions to pedestal formation. Collectively, these results indicate that exocyst-mediated expansion of the plasma membrane acts together with actin polymerization to optimize the generation of pedestals.

**IMPORTANCE:** Enteropathogenic *E. coli* (EPEC) induces the formation of host cell plasma membrane protrusions called “pedestals” that mediate tight adherence to intestinal epithelial cells. Substantial evidence indicates that the generation of pedestals requires bacterial-induced polymerization of the host cell actin cytoskeleton. However, it remains unknown if EPEC manipulates other human processes to augment the efficiency of pedestal generation. The significance of this research is to demonstrate that EPEC co-opts a second host process called polarized exocytosis to increase the frequency of pedestals and the size of these structures. EPEC induces exocytosis by using its effector protein EspH to recruit the human exocyst complex, which mediates the insertion of membrane at sites of bacterial attachment. Our results demonstrate EPEC’s ability to coordinate host membrane trafficking with cytoskeletal changes to enhance infection. Remarkably, EPEC shares this strategy with the intracellular bacterial pathogens *Listeria monocytogenes* and *Shigella flexneri*.

## INTRODUCTION

Enteropathogenic *E. coli* (EPEC) is a major cause of infant diarrhea in developing nations (1–3). EPEC colonizes the apical surface of intestinal epithelial cells by inducing the formation of attaching and effacing (A/E) lesions characterized by the disappearance of microvilli and the production of plasma membrane “pedestals” to which bacteria tightly adhere (1, 3–5). Generation of these pedestals requires bacterial-induced assembly of actin filaments beneath the plasma membrane of the host cell. This actin polymerization is thought to provide a protrusive force that elevates the membrane along with attached bacteria.

Pedestal formation by EPEC requires the bacterium’s type III secretion system (T3SS), which injects ∼ 20 effector proteins into the cytoplasm of human cells (3, 6). One of these effectors is Tir, which inserts into the host plasma membrane and interacts with the EPEC surface protein intimin, resulting in tight adhesion of bacteria (3, 5, 7). The cytoplasmic domain of Tir becomes tyrosine phosphorylated on residue 474 (Y474) by host kinases, resulting in recruitment of the human adaptor protein Nck, which forms a complex with the actin nucleation promoting factor N-WASP (8–10). This latter protein activates the human Arp2/3 complex, thereby stimulating actin polymerization that contributes to the formation of pedestals (8,11). Importantly, pedestal formation promotes bacterial colonization of the intestine (4, 5, 12).

An important unresolved question is whether actin polymerization is the sole host process that contributes to the production of EPEC pedestals. An additional process that could be co-opted by EPEC to reshape the plasma membrane into pedestals is polarized exocytosis: the fusion of intracellular vesicles to expand specific regions in the plasmalemma (13). Polarized exocytosis remodels the plasma membrane of eukaryotic cells during neurite branching, cell migration, phagocytosis, the formation of primary cilia, and other biological events (13–17).

Polarized exocytosis is mediated by the exocyst — a multiprotein complex that is evolutionarily conserved among eukaryotes (13, 18). This complex comprises eight proteins (Sec3, Sec5, Sec6, Sec8, Sec10, Sec15, Exo70 and Exo84) and tethers vesicles to sites in the plasma membrane prior to fusion mediated by SNARE proteins (18, 19). One of the sources of vesicles for the exocyst is the recycling endosome (RE) (13, 18, 19). Fusion of RE-derived vesicles with the plasma membrane is mediated by the v-SNARE protein VAMP3 and various t-SNAREs, including syntaxin 4 (Stx4) (19, 20).

Recent studies reveal that the exocyst complex contributes to the formation of plasma membrane protrusions induced by the bacteria *Listeria monocytogenes* and *Shigella flexneri* (21–23). These protrusions mediate the intercellular spread of *Listeria* or *Shigella* between human cells. Interestingly, *Listeria* and *Shigella* protrusions morphologically resemble EPEC pedestals and are similarly generated by Arp2/3-dependent actin polymerization (5, 24, 25). However, unlike EPEC, *Listeria* and *Shigella* are intracellular pathogens. The actin filaments that promote protrusion formation by these bacteria interface directly with one of the bacterial poles rather than with plasma membrane of the human cell. Nonetheless, the structural similarities between EPEC pedestals and *Listeria* or *Shigella* protrusions prompted us to examine the role of the exocyst in pedestal formation by EPEC.

In this work, we found that EPEC exploits the exocyst to enhance the generation of pedestals. Confocal microscopy imaging showed that polarized exocytosis was locally upregulated in pedestals in a manner dependent on the EPEC T3SS. Of the 15 T3SS effector proteins tested, Tir and the protein EspH made the largest contributions to stimulation of host exocytosis. Depletion of exocyst components by RNA interference (RNAi) or deletion of the *espH* gene each reduced the efficiency of pedestal formation and decreased the size of residual pedestals produced. RNAi-based experiments indicated that the exocyst and Arp2/3 complex work through separate pathways to promote pedestal formation. Taken together, our results reveal an important role for polarized exocytosis in infection by EPEC and demonstrate that manipulation of the exocyst is a strategy that EPEC shares with *Shigella* and *Listeria*.

## RESULTS

### Host exocytosis is stimulated in EPEC pedestals

Exocytosis is augmented in membrane protrusions made by *Listeria* or *Shigella* relative to regions of the host plasma membrane lacking protrusions (21, 22). These findings indicate that these bacteria stimulate polarized exocytosis. To investigate if EPEC induces exocytosis in pedestals, we used an exocytic probe consisting of the v-SNARE protein VAMP3 fused to enhanced green fluorescent protein (EGFP) (14, 21, 26–28). The EGFP part of the probe is inside vesicles and becomes exofacial (extracellular) after fusion of vesicles with the plasma membrane (Fig. 1A). Exocytosis is detected by labeling exofacial VAMP3-EGFP with anti-EGFP antibodies in the absence of membrane-permeabilizing detergents. To determine if exocytosis occurs in EPEC pedestals, we infected human HeLa cells transiently expressing VAMP3-EGFP with the wild-type (WT) EPEC strain E2348/69 for 6 h. Cells were labeled for exofacial EGFP, followed by fixation, permeabilization with a detergent, and labeling of bacteria and filamentous (F)-actin. Images obtained by confocal microscopy demonstrated that exofacial VAMP3-EGFP was present in actin-rich pedestals made by EPEC (Fig. 1B part i). To confirm that exofacial labeling of EGFP was not due to damage-induced permeabilization of the plasma membrane during infection, we performed exofacial labeling on cells that express actin fused to EGFP, which is restricted to the cytosol (21, 22). As expected, intrinsic fluorescence of actin-EGFP was detected regardless of whether infected cells were permeabilized with a detergent (Fig. 1B parts ii and iii). By contrast, the actin-EGFP probe was labeled with EGFP antibodies only when cells were detergent-permeabilized. Collectively, the results in Figure 1B demonstrate that exocytosis occurs in pedestals made by EPEC.

**Figure 1.**
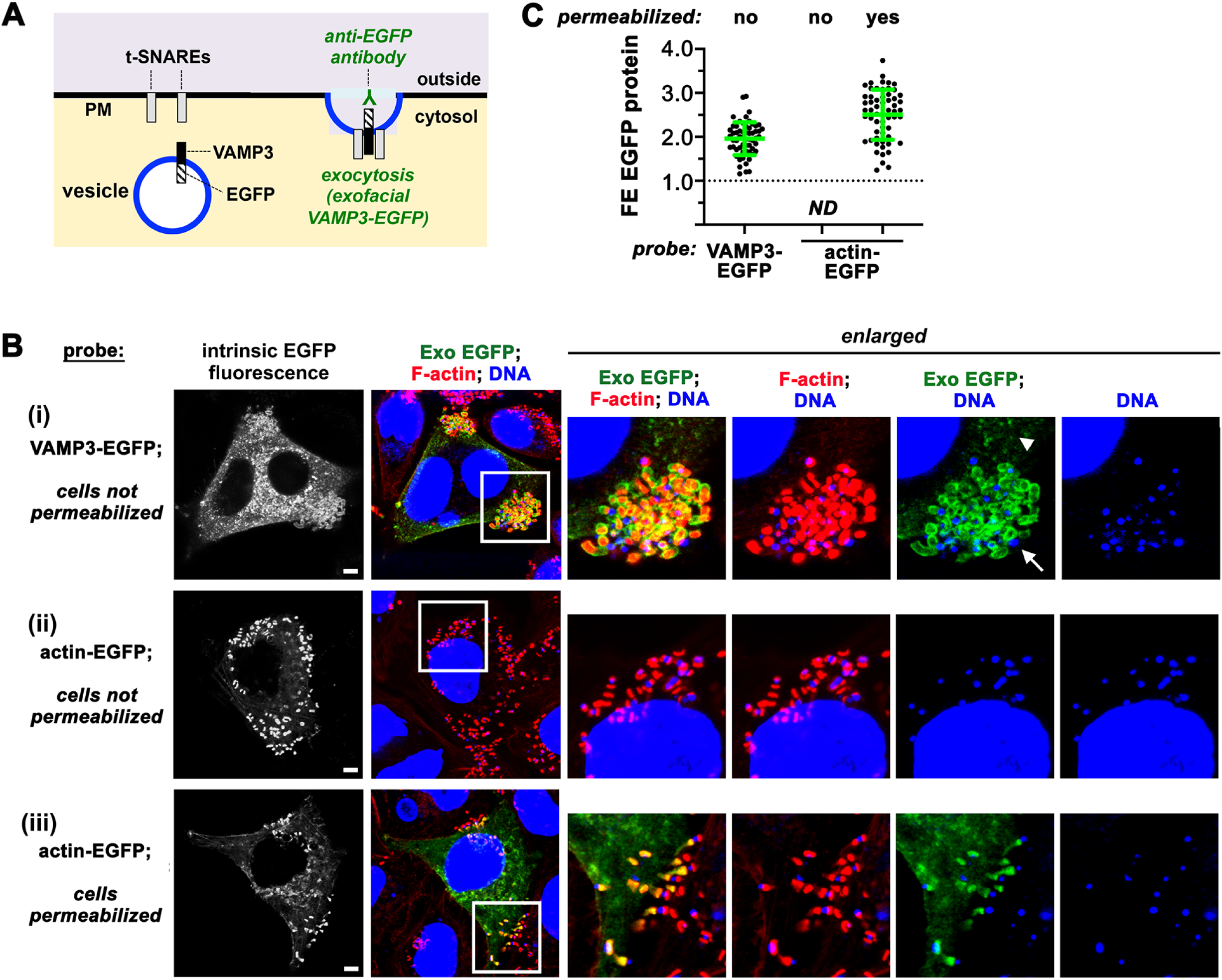
Exocytosis is upregulated in EPEC pedestals. (A). Use of a VAMP3-EGFP probe to assess exocytosis. The EGFP part of VAMP3-EGFP becomes exofacial after vesicles fuse with the plasma membrane. Exocytosis is detected by incubation with unpermeabilized host cells with anti-EGFP antibodies. (B). Representative images of exofacial VAMP3-EGFP (exocytosis) in HeLa cells infected with WT EPEC strain E2348/69. An actin-EGFP fusion protein was used as a negative control for exofacial labeling. The grayscale images show intrinsic fluorescence of the EGFP-tagged proteins, which represents both intracellular and exofacial pools. Scale bars are 5 micrometers. In the remaining images, “Exo EGFP” (green) indicates fluorescence resulting from anti-GFP labeling of VAMP-EGFP or actin-EGFP in unpermeabilized cells. F-actin is colored red. Bacterial and host DNA stained with DAPI are blue. The boxed regions are enlarged views of pedestals and associated exocytosis. (i). Exofacial VAMP3-EGFP in unpermeabilized cells. The arrow indicates a cluster of pedestals and arrowheads show regions of the plasma membrane lacking pedestals. (ii). Lack of exofacial actin-EGFP in unpermeabilized cells. (iii). Detection of actin-EGFP with anti-EGFP antibodies after detergent-permeabilization. (C). The graph displays the degree of accumulation of exofacial VAMP3-EGFP or cytosolic EGFP-actin in pedestals made by EPEC. Data are mean fold enrichment (FE) measurements +/- SD of 59 pedestals for each condition. Each dot is an FE measurement. ND indicates that exofacial EGFP was not detected for actin-EGFP.

To assess if exocytosis is enhanced in pedestals compared to regions of the plasma membrane lacking these structures, we quantified fold enrichment (FE) values, as previously described for *Listeria* and *Shigella* (21, 22). FE is calculated as the mean pixel intensity of exofacial VAMP3-EGFP in bacterial-induced protrusive structures divided by the mean pixel intensity throughout the entire plasma membrane of the infected cell (21, 22). FE values greater than 1.0 indicate an enrichment of exocytosis in protrusive structures. The mean FE value for ∼ 60 pedestals produced by EPEC was 1.95, indicating a nearly two-fold stimulation of exocytosis in these structures (Fig. 1C).

### Exocytosis is dependent on the T3SS of EPEC, Tir-intimin interaction, and the signaling activity of Tir

To investigate the roles of the T3SS and tight adherence mediated by Tir and intimin in induction of exocytosis, HeLa cells expressing VAMP3-EGFP were infected with WT EPEC and isogenic strains deleted for the genes *escN*, *tir,* or *eae*. The *escN* gene encodes a component of a cytoplasmic ATPase complex that associates with the base of the T3SS and is essential for its assembly (29). As a result, the Δ*escN* strain lacks a functional T3SS (30). The *tir* and *eae* genes encode Tir and intimin proteins, respectively. As previously reported, the Δ*escN*, Δ*tir,* and Δ*eae* strains failed to make pedestals on the surface of HeLa cells (7, 30, 31) (Fig. 2A). Nonetheless, we were able to identify bacteria associated with the host cell surface for each of these mutant strains. Measurement of FE values in regions with bound bacteria indicated that the Δ*escN*, Δ*tir*, and Δ*eae* strains were each unable to stimulate exocytosis (Fig. 2B).

**Figure 2.**
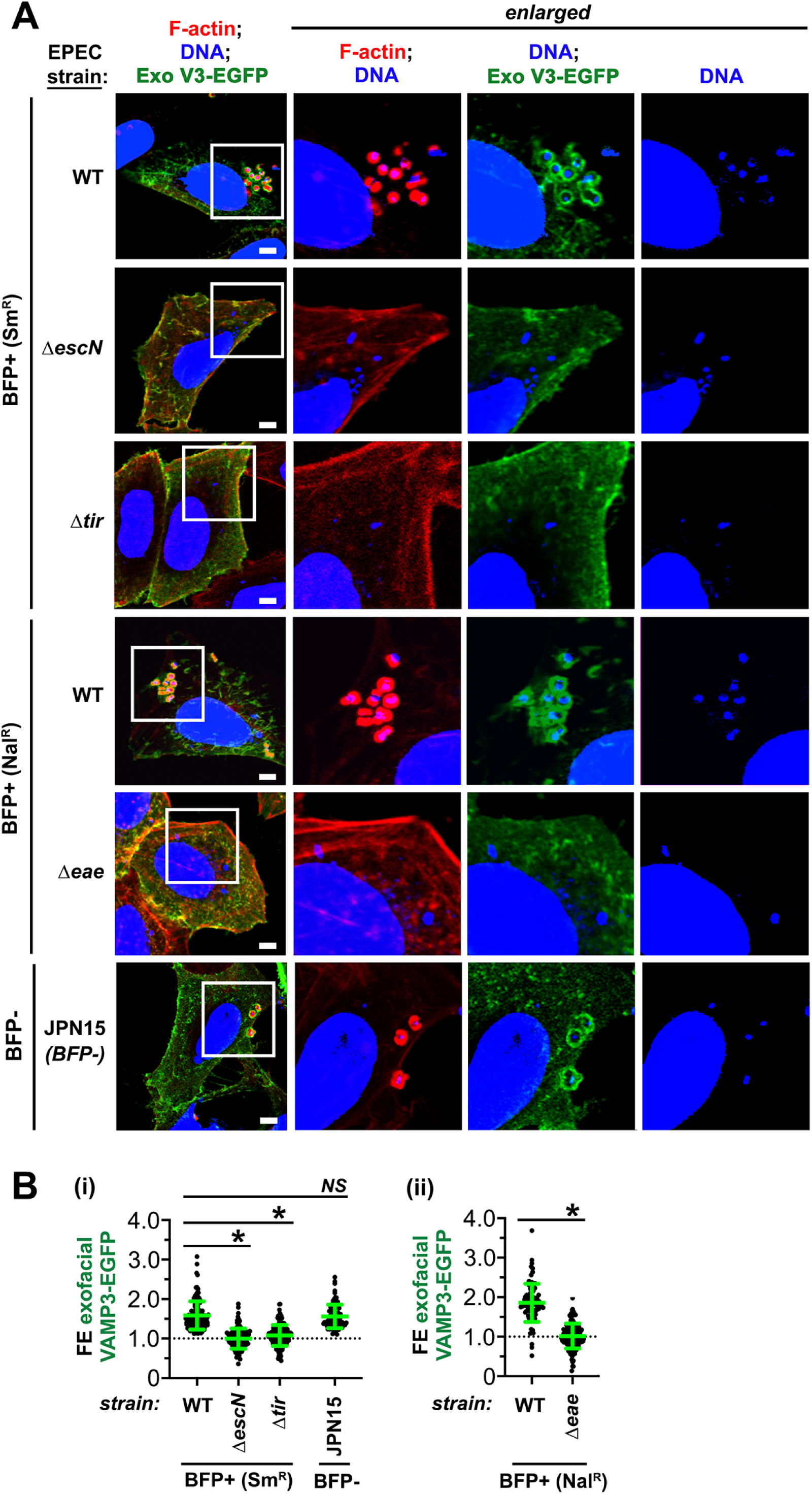
Stimulation of exocytosis by EPEC requires the T3SS, Tir, and intimin. (A). Representative confocal microscopy images of exofacial VAMP3-EGFP (Exo V3-EGFP) in HeLa cells infected for 6 hours with BFP-positive (+) WT streptomycin resistant (Sm^R^) or nalidixic acid (Nal^R^) strains of EPEC, with isogenic mutant strains deleted for *escN*, *tir*, or *eae* genes, or with the BFP-negative (-) strain JPN15. Note the accumulation of exofacial VAMP3-EGFP (Exo V3-EGFP) around pedestals of WT or JPN15 bacteria, but not Δ*escN*, Δ*tir*, or Δ*eae* bacteria. Scale bars are 5 micrometers. The graph shows mean FE values for exofacial VAMP3-EGFP +/- SEM from three experiments. In each experiment, FE measurements of ∼ 40 pedestals were made for each condition. *, P < 0.05. NS indicates not significant.

EPEC has a type IV “bundle forming” pilus (BFP), which binds to host cells prior to secretion by the T3SS and subsequent tight adherence mediated by intimin and Tir (6, 32). To assess the role of the BFP in upregulation of exocytosis, we used EPEC strain JPN15, which is a derivative of E2348/69 that lacks a virulence plasmid encoding the BFP (33, 34). JPN15 (BFP-) was found to stimulate exocytosis as efficiently as WT (BFP+) EPEC (Fig. 2), revealing that the BFP is dispensable for induction of exocytosis.

Phosphorylation of Tir on tyrosine 474 stimulates a host signaling pathway that activates the Arp2/3 complex, resulting in actin polymerization in pedestals (3, 8, 9). To assess if the signaling function of Tir is needed for upregulation of host exocytosis, JPN15-derived strains deleted for the *tir* gene (Δ*tir*) and expressing plasmid-borne wild-type Tir protein or Tir with a tyrosine-to-phenylalanine substitution at residue 474 (Y474F) were used. The results indicate that the strain expressing Tyr474F was unable to support induction of exocytosis (Fig. S1). Taken together, findings in Figures 2 and S1 show that the ability of EPEC to stimulate exocytosis depends on the T3SS, intimate adherence, and Tir-mediated signaling, but not on the BFP.

### The host exocyst complex contributes to the formation of EPEC pedestals

Polarized exocytosis in protrusions of *Listeria* or *Shigella* is mediated by the human exocyst (21, 22). Experiments involving RNA interference (RNAi)-mediated depletion of exocyst components indicate that the exocyst is needed for the efficient formation of protrusions by these bacteria. We performed similar RNAi experiments to assess the role of the exocyst in pedestal formation by EPEC. HeLa cells were transfected with short interfering RNAs (siRNAs) targeting the exocyst proteins Sec3, Sec6, Sec8, Sec10, or Exo70 (Fig. 3A). The v-SNARE protein VAMP3 and the t-SNARE syntaxin 4 (Stx4) were also targeted with siRNAs. As negative controls, cells were mock transfected in the absence of siRNA or transfected with a control siRNA that lacks complementarity to any known human mRNA. As a positive control expected to impair pedestal formation, cells were transfected with an siRNA against the Arp3 component of the Arp2/3 complex. After transfection, HeLa cells were infected with EPEC strain JPN15 for 6 hours, followed by fixation, permeabilization with a detergent, and labeling of bacteria and F-actin in pedestals. In parallel with these infection experiments, Western blotting studies were performed to demonstrate effective depletion of exocyst components, VAMP3, and Stx4 by siRNAs (Fig. S2).

**Figure 3.**
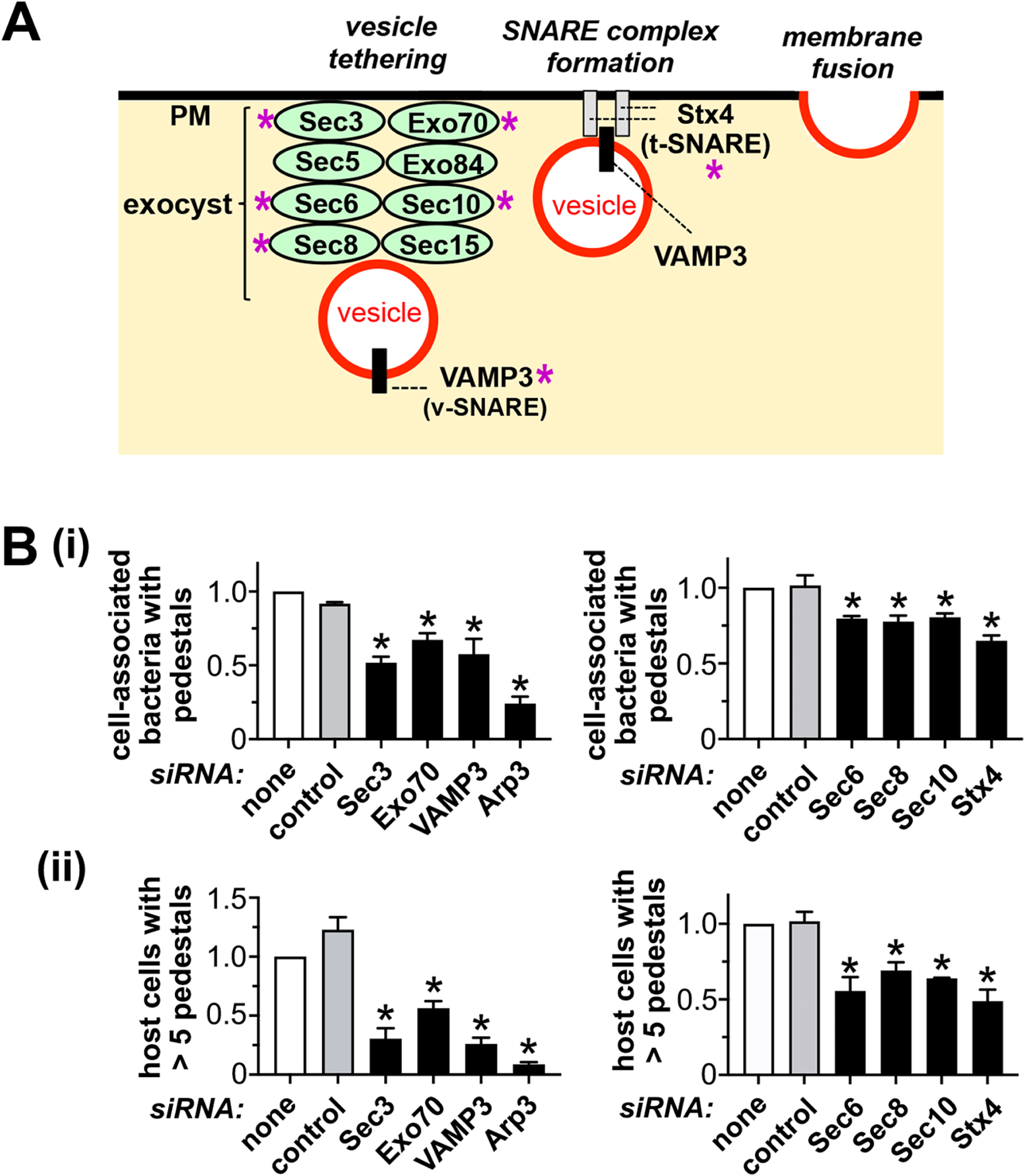
The exocyst and the SNARE proteins VAMP3 contribute to pedestal formation by EPEC. (A). The exocyst complex tethers vesicles to sites in the plasma membrane (PM) followed by membrane fusion mediated by the SNARE proteins VAMP3 and Stx4 (13, 18, 19). Proteins indicated with asterisks were targeted by RNAi. (B). Effect of siRNAs against exocyst proteins, VAMP3, or Stx4 on pedestal formation by strain JPN15 in HeLa cells. Pedestal formation efficiency was expressed as the relative percentage of cell-associated bacteria that make pedestals (i) or the relative percentage of HeLa cells with more than 5 pedestals (ii). The data are mean relative pedestal formation values +/- SEM from 3-4 experiments, depending on the condition. *, P < 0.05 compared to the control siRNA condition.

Pedestal formation was assessed by confocal microscopy imaging. Two methods were used to quantify the efficiency of pedestal generation: the percentage of cell-associated bacteria that have pedestals (35, 36) and the percentage of infected cells that have 5 or more pedestals (37). We reasoned that the second method might provide a more sensitive means of detecting defects in pedestal production. The rationale for this idea is that highly infected cells may be limited in plasma membrane capacity and therefore more reliant on exocyst-mediated insertion of membrane for pedestal formation. The results from the RNAi studies indicate that siRNA-induced depletion of exocyst components, VAMP3, or Stx4 caused a ∼ 20-50% decrease in the percentage of cell-associated bacteria that made pedestals (Fig. 3B part i) and a ∼ 30-80% reduction in the proportion of host cells with more than 5 pedestals (Fig. 3B part ii). As expected, depletion of Arp3 also impaired pedestal formation. Taken together, these findings indicate that exocyst components and SNARE proteins are needed for the efficient production of pedestals.

### The exocyst is recruited to sites of pedestal formation and mediates exocytosis in these structures

Using an EGFP-tagged probe, we found that the exocyst component Exo70 is enriched in pedestals made by WT JPN15, but does not accumulate around Δ*tir* mutant bacteria associated with the HeLa cell surface (Fig. 4). Importantly, siRNA-mediated depletion of exocyst components or the SNARE protein Stx4 reduced enrichment of exocytosis in JPN15 pedestals (Fig. 5). Taken together, the results in Figures 4 and 5 indicate that EPEC manipulates the exocyst by mobilizing the complex to protrusion sites, where it promotes exocytosis.

**Figure 4.**
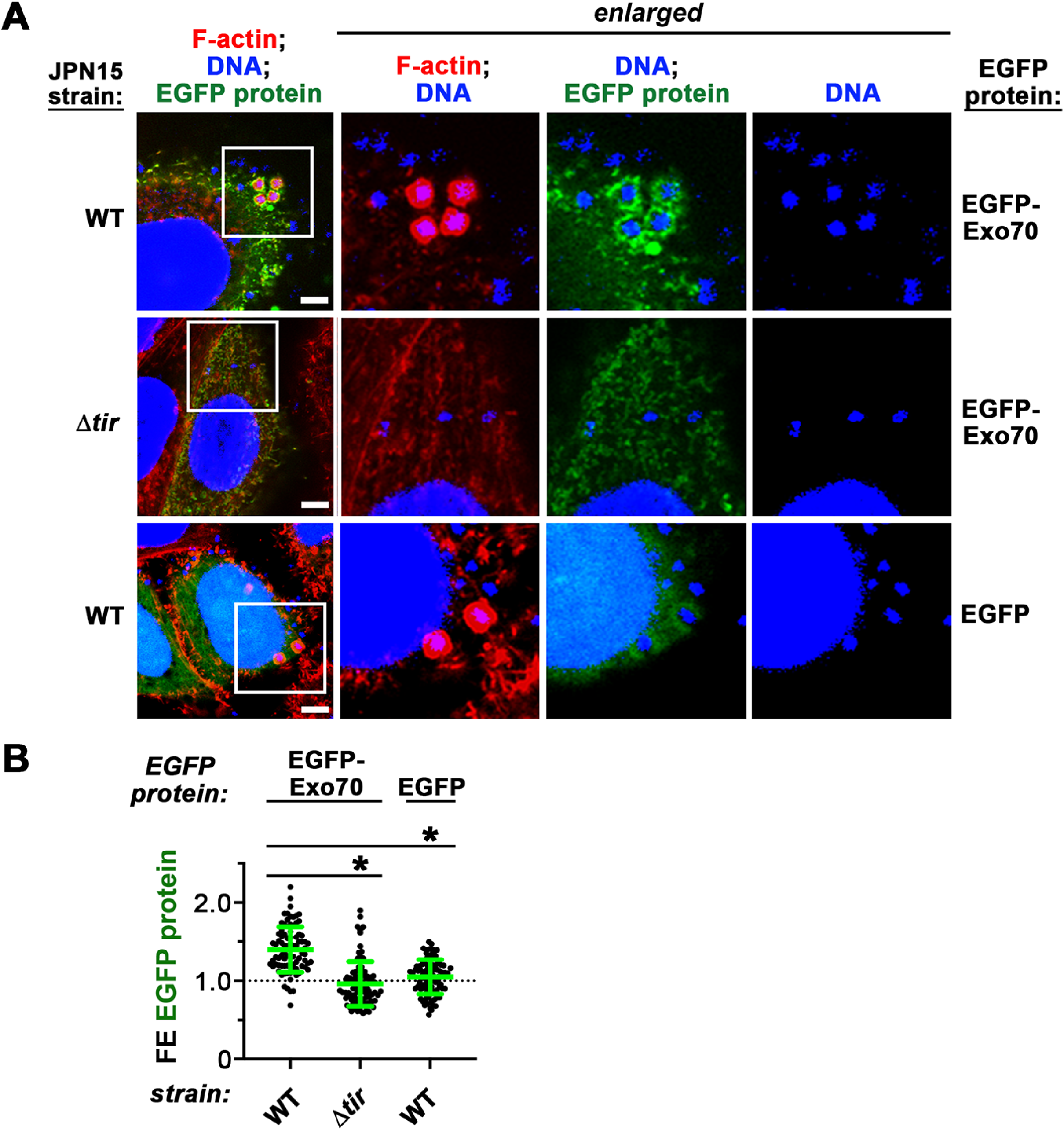
The exocyst component Exo70 is recruited to EPEC pedestals. HeLa cells transiently expressing EGFP-tagged Exo70, EGFP-TC10, or EGFP alone were infected with JPN15 or JPN15 deleted for *tir* (Δ*tir*) as a control for 6 h, followed by fixation, permeabilization, and labeling for F-actin in pedestals or DAPI to stain host and bacterial DNA. (A). Confocal microscopy images showing accumulation of EGFP-Exo70 around pedestals. (B). A graph with mean FE values +/- SD from three experiments is presented. Approximately 30 pedestals were measured for each condition in each experiment. *, P < 0.05.

**Figure 5.**
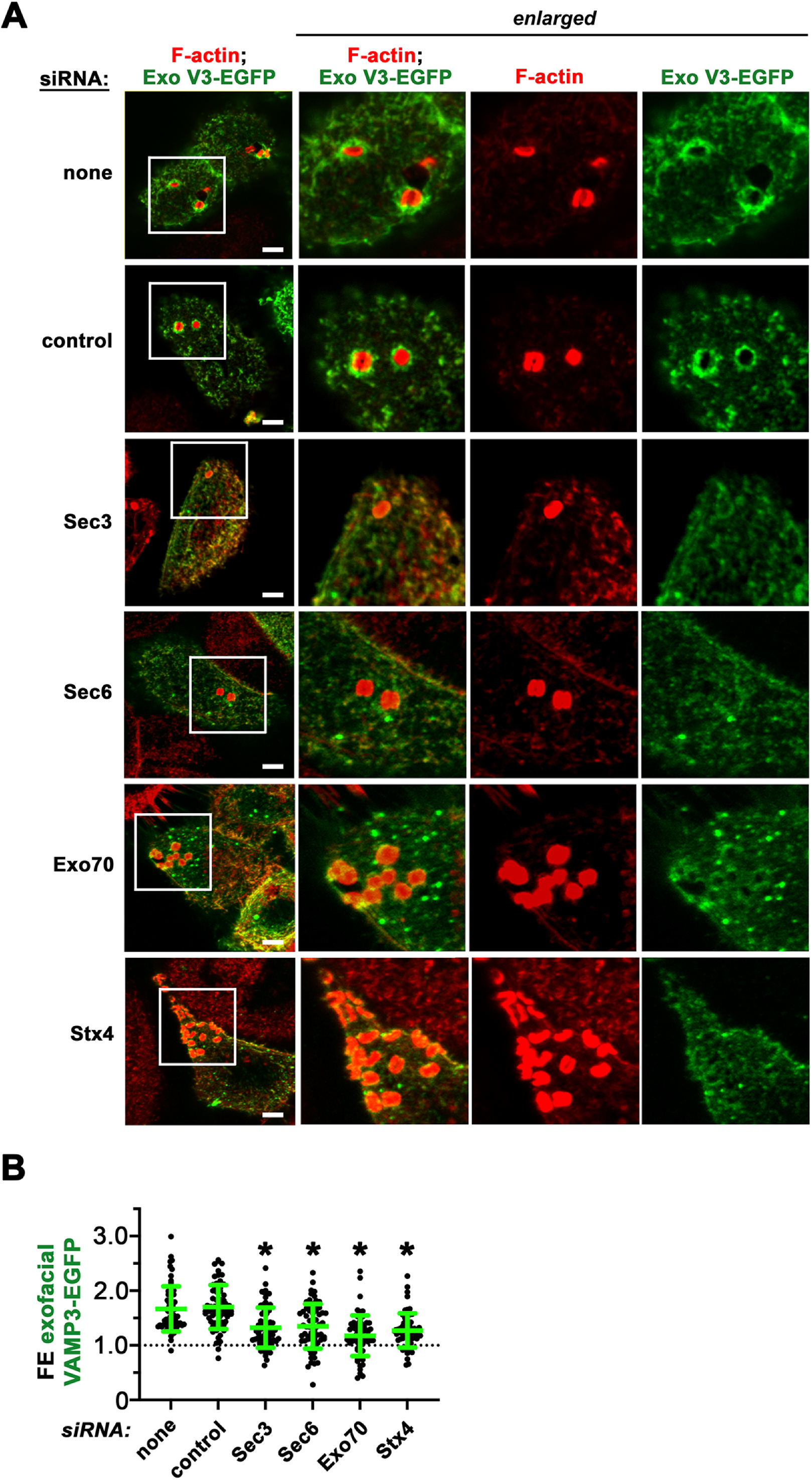
The exocyst complex mediates exocytosis in EPEC pedestals. HeLa cells were either mock transfected in the absence of siRNA (none), transfected with a control non-targeting siRNA, or transfected with siRNAs against Sec3, Sec6, Exo70, or Stx4. Cells were then transfected with a plasmid expressing VAMP3-EGFP, infected with the EPEC strain JPN15 for 6 h and labeled for exofacial VAMP3-EGFP, F-actin, and bacteria as described in the Materials and Methods. (A). Representative confocal microscopy images of EPEC pedestals are presented. Exofacial VAMP3-EGFP (Exo V3-EGFP) is green and F-actin is red. (B). The graph shows mean FE values +/- SD for exofacial VAMP3-EGFP in pedestals from three experiments. In each experiment, approximately 25 pedestals were measured for each condition. *, P < 0.05 compared to the no siRNA or control siRNA conditions.

### The exocyst and Arp2/3 complex act separately to promote pedestal formation

Components of the exocyst are known to interact with the Arp2/3 complex or members of the WAVE regulatory complex, indicating that the exocyst has the potential to control actin filament assembly (38, 39). We therefore investigated if actin polymerization in EPEC pedestals depends on the exocyst complex. RNAi-mediated depletion of exocyst or SNARE proteins failed to reduce enrichment of F-actin in pedestals of JPN15 (Fig. 6). Instead, siRNAs targeting Sec6, Sec8, Sec10, Exo70, VAMP3, or Stx4 caused 7-20% mean increases in F-actin accumulation in pedestals. These findings indicate that the exocyst is dispensable for actin filament assembly in pedestals.

**Figure 6.**
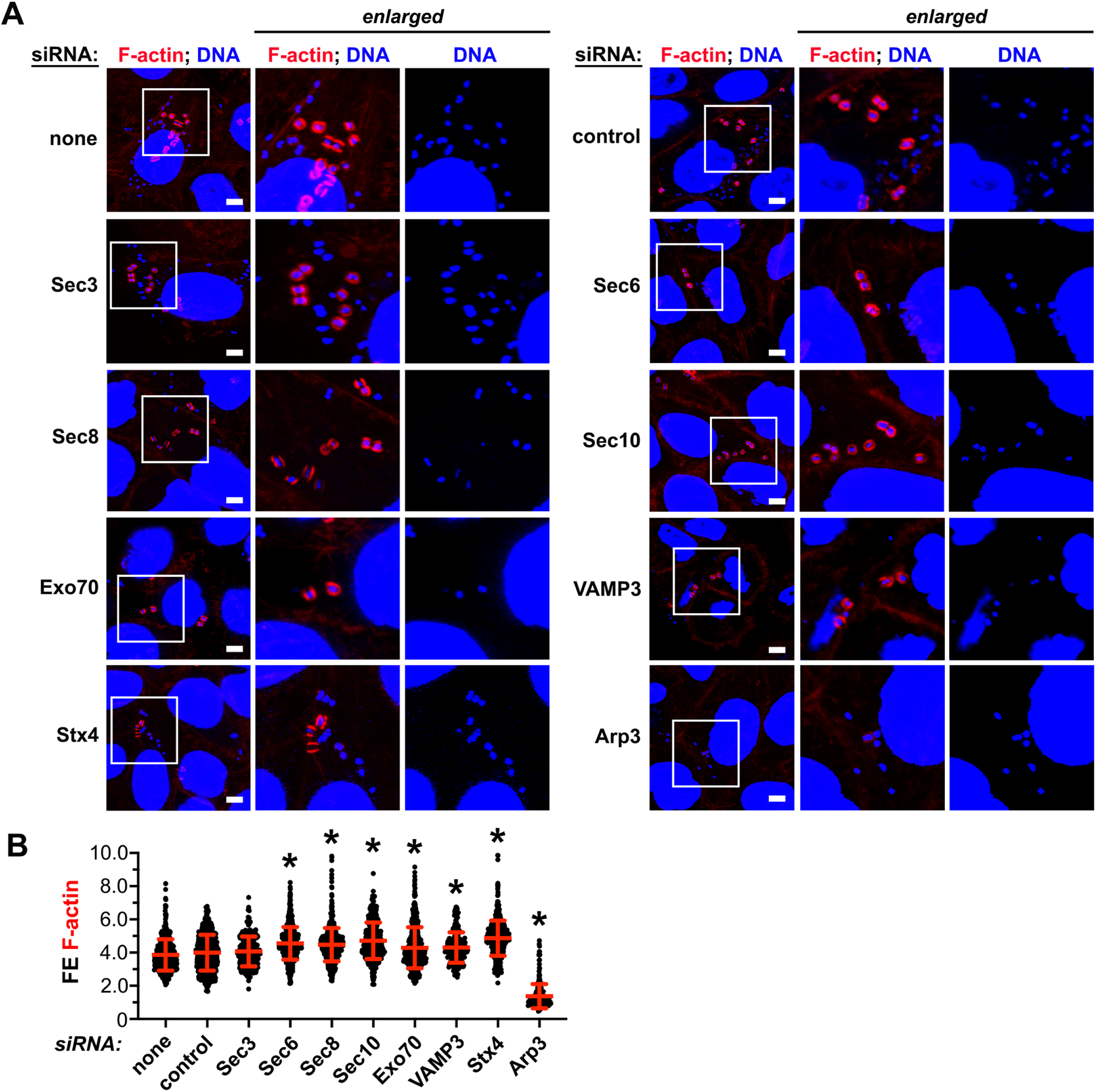
Exocyst and SNARE proteins are dispensable for accumulation of filamentous (F)-actin in pedestals. HeLa cells were mock transfected in the absence of siRNA, transfected with a control siRNA, or transfected with siRNAs targeting Sec3, Sec6, Sec8, Sec10, Exo70, VAMP3, or Stx4. As a control expected to decrease F-actin in pedestals, cells were transfected with an siRNA against Arp3. HeLa cells were then infected with EPEC strain JPN15 for 6 h, followed by fixation, permeabilization, and labeling of F-actin and DNA. (A). Shown are representative confocal microscopy images with F-actin in red and DNA in blue. Scale bars represent 5 micrometers. (B). Enrichment of F-actin in pedestals. The graph shows mean FE values +/- SD in pedestals from three experiments. Approximately 120-200 pedestals were analyzed for F-actin enrichment in each experiment. *, P < 0.05. NS indicates not significant.

The relationship between exocytosis and actin polymerization was further investigated by examining the impact of co-depletion of exocyst components and Arp3 on pedestal formation efficiency. When HeLa cells were co-depleted for Sec3, Exo70, or VAMP3 along with Arp3, protrusion formation decreased relative to depletion of Arp3 alone (Fig. 7). Collectively, the findings in Figures 6 and 7 provide evidence that exocytosis and actin polymerization act on separate pathways to control pedestal formation.

**Figure 7.**
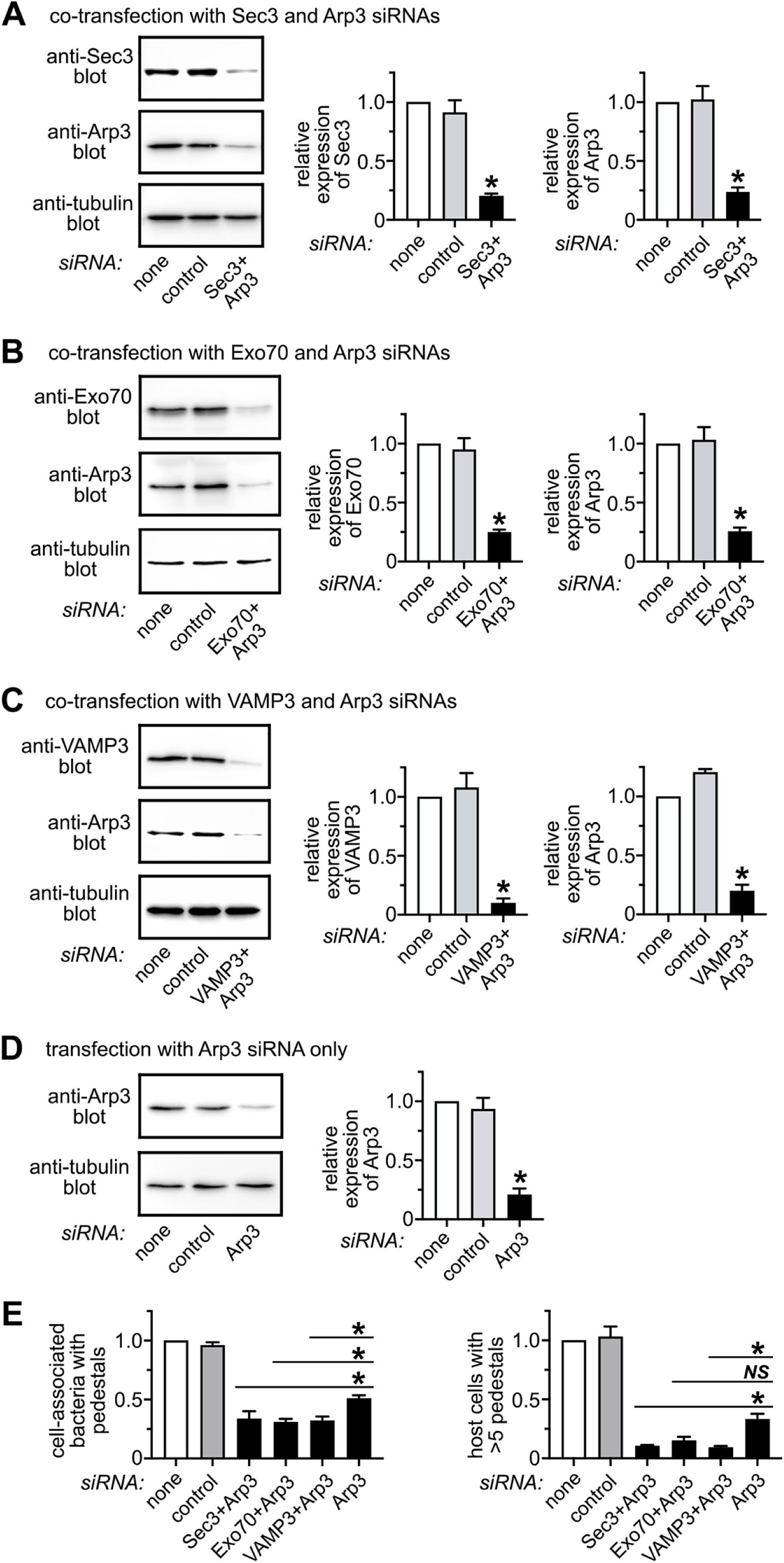
Proteins involved in exocytosis or actin polymerization contribute additively to pedestal formation. HeLa cells were subjected to control conditions or co-transfected with siRNAs targeting Arp3 and Sec3, Arp3 and Exo70, or Arp3 and VAMP3. Cells were infected with strain JPN15 for 6 h, followed by fixation, permeabilization, and labeling for F-actin in pedestals and with DAPI to detect bacteria. (A, B, C, and D). Representative Western blots confirming depletion of Sec3, Exo70, VAMP3, or Arp3 are presented. Data are mean +/- relative expression values from three experiments. (E) Mean relative protrusion efficiencies +/- SD from three experiments are shown. *, P <0.05. NS indicates not significant.

### Identification of EPEC effector proteins that promote host exocytosis

The finding that the T3SS is needed for upregulation of exocytosis in pedestals (Fig. 2) suggested that one or more EPEC effector contributes to induction of exocytosis. Our results indicate that one such effector is Tir (Figures 2 and S1). To identify other effectors that may promote exocytosis, we screened BFP-positive strains of EPEC that were deleted for 14 different effector genes (Figures 8 and S3). Relative to isogenic WT strains, mutant strains deleted for *espH*, *map*, *nleA*, or *espF* genes were partly defective in stimulation of exocytosis in pedestals (Figure 8). Of these four mutant strains, the Δ*espH* strain exhibited the greatest defect in enhancement of exocytosis. The remaining 10 effector mutant strains induced exocytosis similarly to WT strains (Figures 8B and S3).

**Figure 8.**
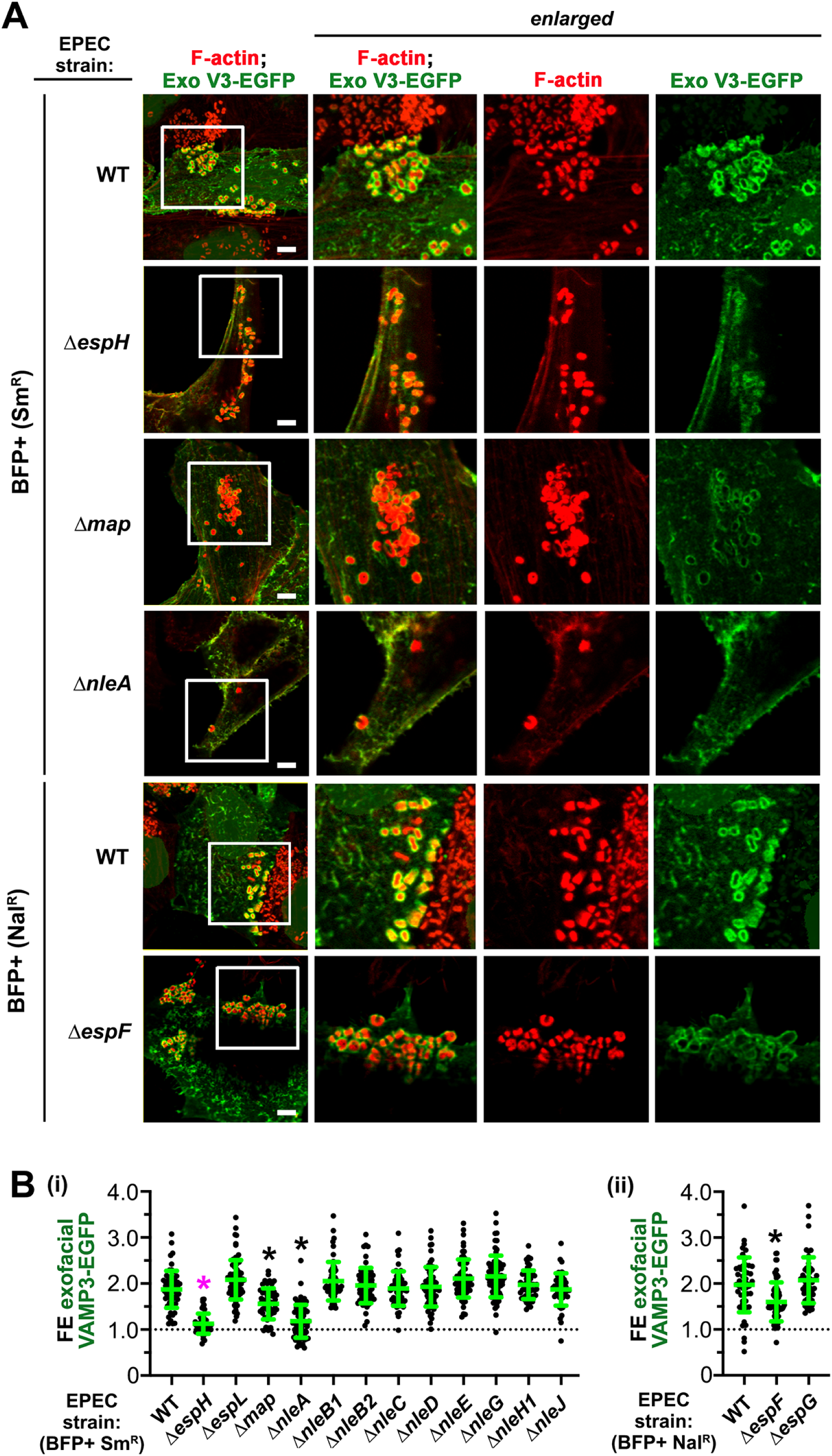
Identification of T3SS effectors that contribute to stimulation of exocytosis. HeLa cells transiently expressing VAMP3-EGFP were infected for 6 h with BFP-positive (+) Sm^R^ or Nal^R^ strains of WT EPEC or with isogenic mutant strains deleted for genes encoding specific effector proteins. Cells were then permeabilized and labeled for F-actin to detect pedestals. (A). Shown are representative confocal microscopy images of exofacial VAMP3-EGFP (Exo V3- EGFP) in pedestals made by WT EPEC or mutant strains that exhibited decreased ability to induce exocytosis. (B). The graphs in i and ii contain exocytosis results from all 14 mutant strains tested. The Δ*espH* strain is highlighted with a magenta asterisk. The data are mean FE values for exofacial VAMP3-EGFP +/- SD from 52-83 pedestals, depending on the condition. *, P <0.05, compared to the mean FE value of the WT Sm^R^ or Nal^R^ strains.

### EspH mobilizes the exocyst to promote exocytosis and pedestal formation

Using a JPN15-derived strain deleted for the *espH* gene (40), we confirmed that EspH contributes to induction of exocytosis by BFP-negative bacteria (Figure 9A). Importantly, the Δ*espH* mutant strain also exhibited reduced recruitment of EGFP-tagged Exo70 (Figure 9B) or endogenous Exo70 (Figure S4) and a lower efficiency of pedestal formation compared to the wild-type strain (Figure 9C). The defects of the Δ*espH* strain in stimulation of exocytosis, recruitment of EGFP-Exo70, and pedestal formation were complemented by expression of a functional *espH* gene on a plasmid (Figure S5). Taken together, the results in Figures 9, S4, and S5 provide evidence that EspH promotes exocytosis and pedestal formation and that these activities are linked to the effector’s ability to alter the subcellular localization of Exo70.

**Figure 9.**
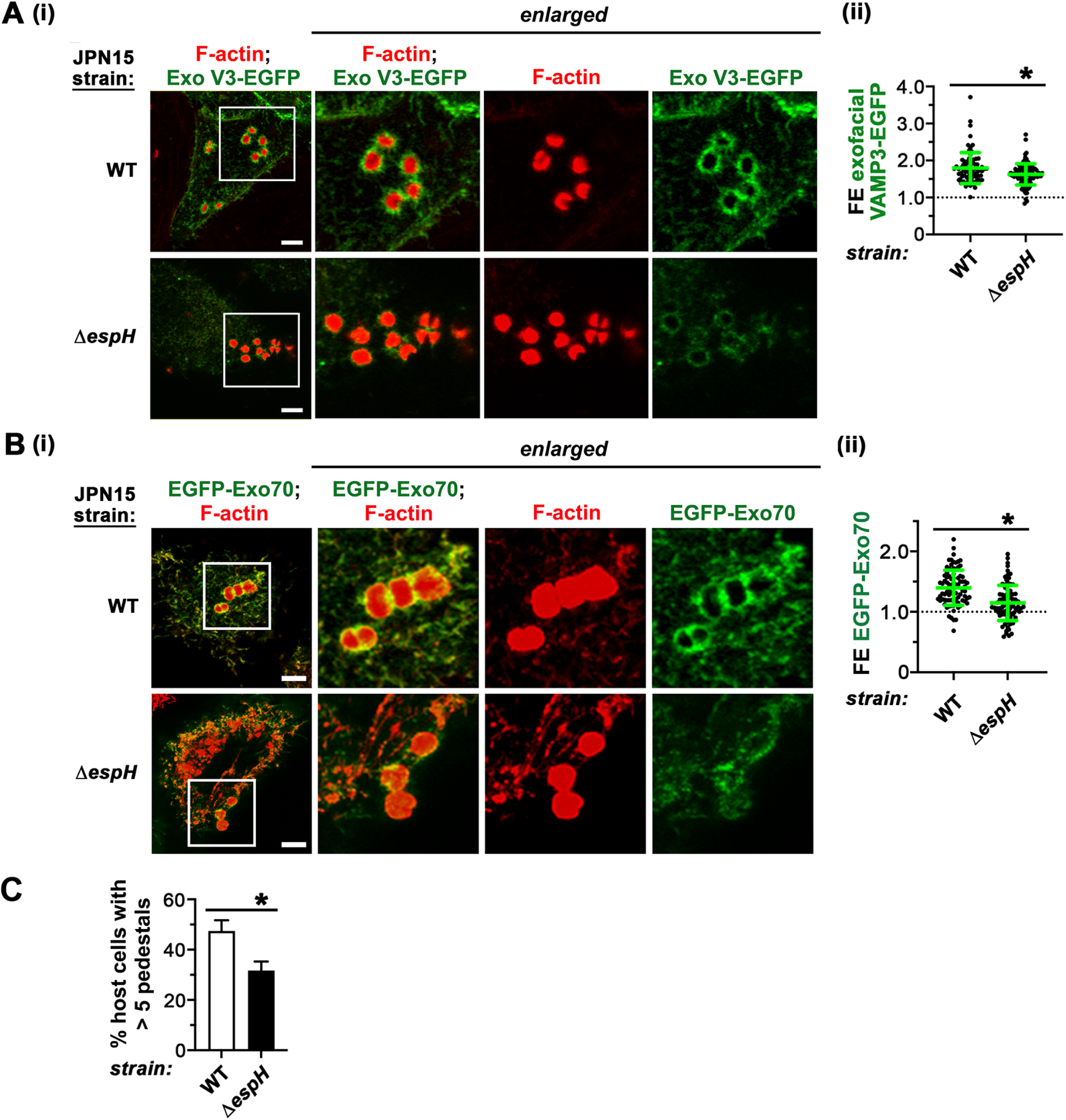
EspH promotes exocytosis, recruitment of Exo70, and formation of pedestals. HeLa cells transiently expressing VAMP3-EGFP or EGFP-Exo70 were infected with WT JPN15 or an isogenic strain deleted for the *espH* gene (Δ*espH*) for 6 h, followed by fixation and labeling for exofacial VAMP3-EGFP and F-actin in pedestals. Results for enrichment of exocytosis or EGFP-Exo70 in pedestals are presented in parts A, B, respectively. Data in the graphs are mean FE values +/- SD from three experiments. In each experiment, 25-35 pedestals were assessed for each condition. (C). Comparison of pedestal formation efficiencies between WT and Δ*espH* strains of JPN15. The data are from three experiments. *, P < 0.05.

### EspH affects the spatial distribution of exocytosis and Exo70 around pedestals

Deletion of the *espH* gene reduces, but does not abolish, exocytosis and localization of Exo70 around pedestals (Figures 8, 9, and S4). To gain insight on the effects of EspH on the spatial distribution of exocytosis during infection, we assembled 60 confocal microscopy images taken at 230 nm intervals in the Z plane to reconstruct a 3-dimensional (3D) view of exocytosis associated with pedestals. The results reveal that exocytosis is contiguous around pedestals of WT JPN15, whereas the Δ*espH* mutant strain exhibits gaps in pedestal-associated exocytosis (Fig. 10A; supplemental movies 1 and 2). Similar gaps in localization of EGFP-Exo70 were observed around pedestals made by the Δ*espH* mutant strain (Fig. S6A; supplemental movies 3 and 4). These results indicate that EspH supports even distribution of exocytosis around pedestals, in addition to enhancing the overall level of exocytosis in these structures.

**Fig. 10.**
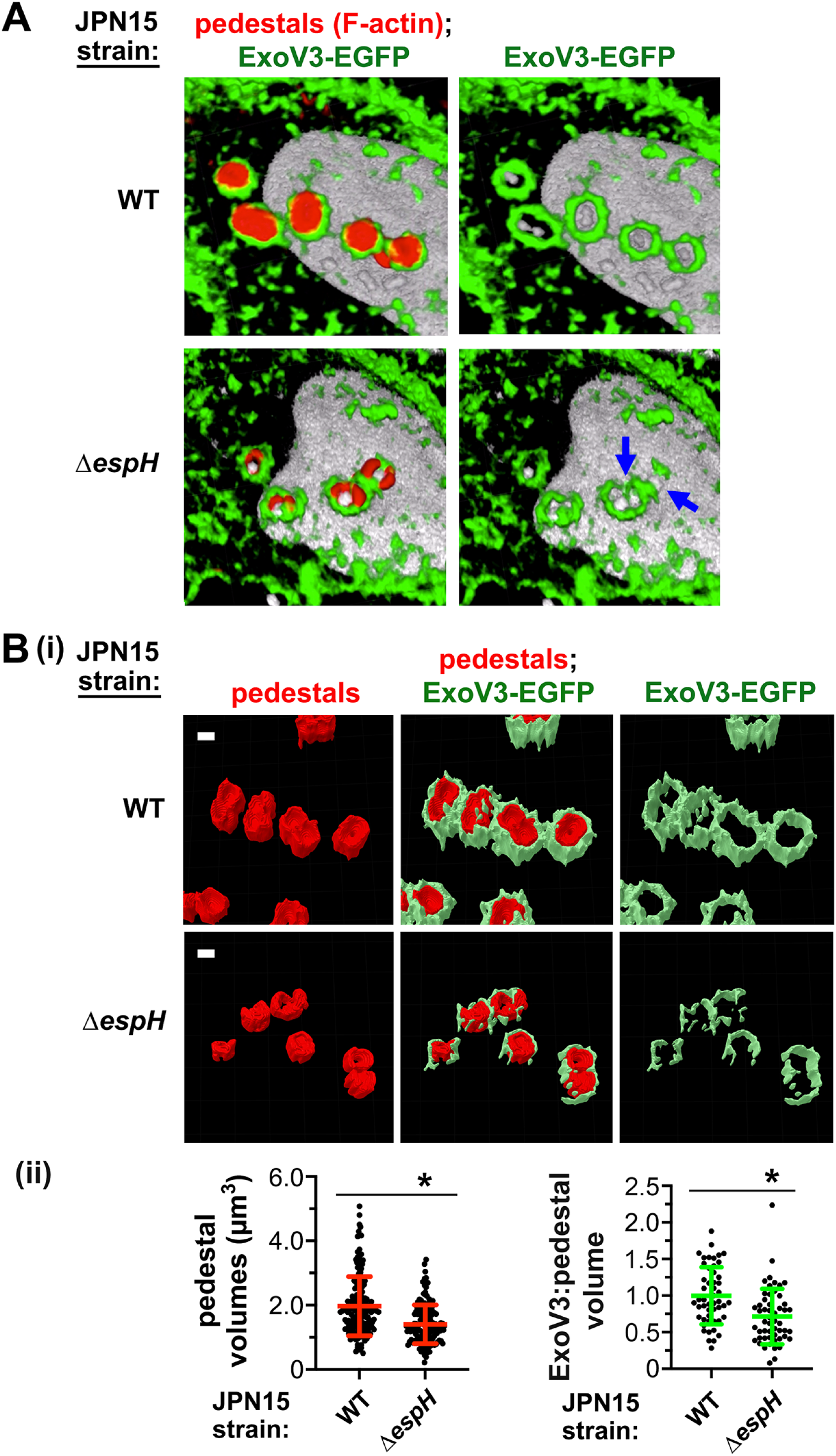
EspH controls the spatial distribution of exocytosis and the size of pedestals. HeLa cells transiently expressing VAMP3-EGFP were infected with WT or Δ*espH* strains of JPN15 for 6 h. Cells with VAMP3-EGFP were subjected to exofacial labeling with anti-EGFP antibodies and then fixed, permeabilized, and labeled for F-actin and DNA. Part A is a three-dimensional (3D) reconstruction of Z stack images of exocytosis (Exo V3-EGFP) around pedestals of WT and Δ*espH* strains. Arrows indicate gaps in exocytosis around pedestals of the Δ*espH* strain. These data are also shown in supplementary movies 1 and 2. Part B contains images of pedestals and exocytosis obtained from Airyscan confocal microscopy images processed using Zeiss arivis Pro software. Scale bars are 1 micrometer. The graphs in part C display pedestal volumes (i) or the ratio of exocytosis (ExoV3) to pedestal volumes (ii). Horizontal bars indicate means and vertical bars are SD. For part (i), volumes of 163 pedestals of WT JPN15 and 152 pedestals of the Δ*espH* strain were quantified. For part (ii), volumes of ExoV3-EGFP and pedestals were measured for 50 and 53 pedestals of WT or Δ*espH* strains, respectively. *, P < 0.05.

### EspH and the exocyst control the size of EPEC pedestals

To examine the impact of EspH on the dimensions of pedestals, Airyscan confocal microscopy (41) was used to obtain super resolution images of pedestals and associated exocytosis produced by WT and Δ*espH* strains of BFP-negative JPN15. These images were then refined by processing with arivis Pro software. The volumes of ∼ 150 pedestals were quantified, revealing that the mean pedestal size of the Δ*espH* strain was ∼ 30% smaller than that of the WT strain (Fig. 10B). In addition, the Δ*espH* strain had reduced volumes of exocytosis surrounding pedestals. Similar results were obtained for the volumes of EGFP-tagged Exo70 around pedestals (Fig. S6B). We also examined whether EspH controls the size of pedestals of BFP-positive bacteria. The BFP clusters bacteria into groups, making it difficult to analyze individual pedestals with confocal microscopy. We therefore used serial block face scanning electron microscopy (SBF-SEM) to acquire images of individual pedestals of BFP- positive bacteria. Measurements of the lengths and areas of these pedestals indicated that the BFP-positive Δ*espH* mutant strain makes smaller pedestals compared to BFP- positive WT bacteria (Fig. S7). Finally, we examined the role of the exocyst and SNARE proteins in controlling the size of pedestals made by WT JPN15 (BFP-) bacteria. Targeting the exocyst component Sec3 or the SNARE protein Stx4 reduced mean pedestal volumes compared to the control siRNA condition by 30 or 23%, respectively (Fig. 11). Taken together, the findings in Figures 10B, 11, S6B, and S7 provide evidence that EspH and host proteins that mediate exocytosis promote the expansion of pedestals.

**Figure 11.**
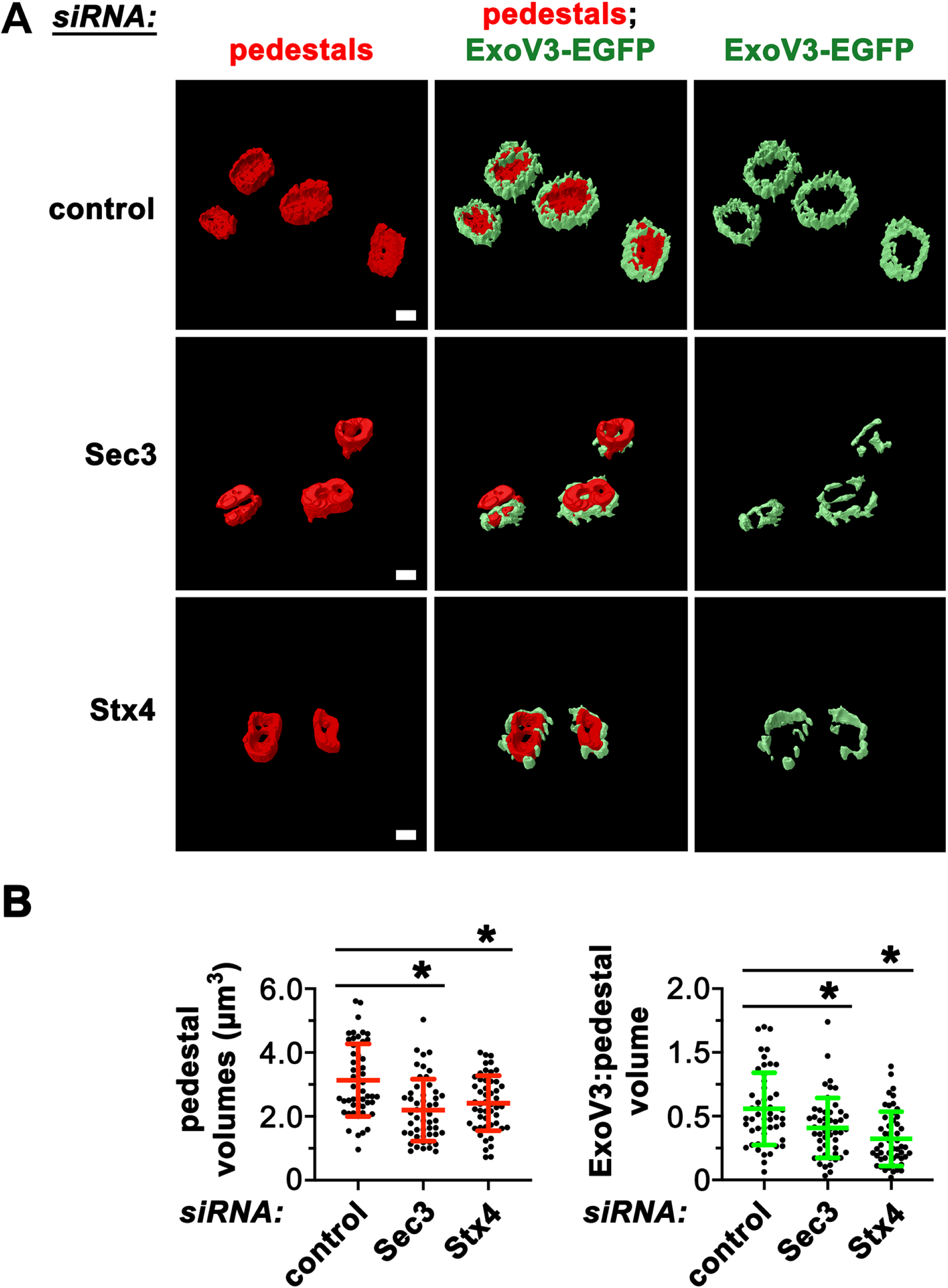
Host proteins involved in exocytosis regulate pedestal size. HeLa cells transiently expressing VAMP3-EGFP were subjected to control conditions or transfected with siRNAs targeting Sec3 or VAMP3 for approximately 48 h. Cells were then infected with WT or Δ*espH* strains of JPN15 for 6 h, followed by fixation, permeabilization, and labeling for F-actin and DNA. Samples were imaged by Airyscan confocal microscopy and processed using Zeiss arivis Pro software. (A). Representative images of pedestals are presented. Scale bars are 1 micrometer. (B). Data in graphs are mean volumes +/- SD for pedestals. Approximately 75 pedestals were analyzed for each of the three conditions. *, P < 0.05.

## DISCUSSION

In this work, we demonstrated that the human exocyst complex controls the frequency of formation of EPEC pedestals, as well as the size of these structures. EPEC manipulates the exocyst, at least in part, through its effector protein EspH, which promotes recruitment of the exocyst component Exo70. In addition, our studies provide evidence that exocytosis mediated by the exocyst and actin polymerization induced by the Arp2/3 complex act separately to contribute to pedestal formation.

Our experiments involved detection of exocytosis using a VAMP3-EGFP fusion protein (14, 21, 22, 26–28). Since VAMP3 localizes in vesicles from the recycling endosome (RE) (19, 20), the VAMP3-EGFP probe detects exocytosis of RE-derived membrane. It was previously reported that host proteins that control RE- mediated recycling, including VAMP3, Exo70, and the GTPase Rab11a are recruited to sites of EPEC attachment to human cells (42, 43). However, these studies did not demonstrate exocytosis of RE-derived membrane in pedestals, nor did they investigate the roles of the exocyst or VAMP3 in pedestal formation and colonization of host cells.

Our finding that the T3SS effector EspH enhances the efficiency of pedestal formation is in agreement with results from two previous studies (40, 44). Although these studies also reported that EspH affects the lengths of pedestals, the data provided were solely visual and not supported by measurements of pedestal dimensions. Our study used Airyscan confocal microscopy and SBF-SEM to measure the volumes, lengths, or areas of pedestals produced by WT and Δ*espH* mutant bacteria, thereby definitively demonstrating that EspH controls pedestal size. In addition, by showing that EspH is needed for stimulation of exocytosis and recruitment of Exo70, our results provide mechanistic information for how EspH affects the formation and size of pedestals.

Apart from EspH, our study identified potential roles for EspF, Map, and NleA in stimulation of exocytosis. In general agreement with these findings, EspF and Map were previously found to promote the recruitment of host proteins in RE-derived vesicles to sites of EPEC adherence (42). Map is an activator of the human GTPase Cdc42, which is known to positively regulate the exocyst (13, 18, 45). It is therefore possible that Map controls exocytosis via manipulation of Cdc42. The mechanism by which EspF contributes to EPEC-induced exocytosis remains unknown but could involve its host ligand SNX9 (46). The effector protein NleA binds to the COPII coat component Sec24, thereby inhibiting vesicle transport from the endoplasmic reticulum (ER) to the Golgi apparatus (47). The exocyst can mediate exocytosis using not only RE-derived vesicles, but also vesicles that bud from the *trans*-Golgi network (13, 18). However, it is unclear how blocking ER-Golgi transport by NleA could enhance exocytosis in EPEC pedestals. Indeed, one would expect inhibition of Sec24 by NleA to impair exocytosis. We suggest that NleA may target unidentified host ligands to manipulate the exocyst.

In addition to stimulating RE-dependent exocytosis, EPEC has been found to induce the separate exocytic pathway of lysosomal exocytosis (48, 49). This pathway involves the fusion of lysosomes with the plasma membrane and is independent of most components of the exocyst and of the v-SNARE protein VAMP3 (50, 51). Instead, fusion of lysosomes with the plasma membrane is mediated by the v-SNARE protein VAMP7 (52). Interestingly, the ability of EPEC to induce lysosomal exocytosis depends on EspH (48, 49). Whether lysosomal exocytosis affects pedestal formation and/or size has not been reported.

In summary, our findings demonstrate an important role for the human exocyst complex in pedestal generation by EPEC. Remarkably, EPEC shares with the bacteria *Listeria* and *Shigella* the ability to manipulate the exocyst to enhance the production of protrusive structures that contribute to infection (21, 22). Future work should reveal mechanisms by which virulence proteins of these pathogens control exocyst function and how exocytosis is spatially and temporally coordinated with actin polymerization to optimize protrusion formation efficiency.

## ACKNOWLEDGEMENTS

We thank Dr. Ilan Roshenshine for wild-type and Δ*espH* strains of JPN15. Dr. Julian Guttman is gratefully acknowledged for advice on quantification of pedestal formation. Some of the imaging in this study was performed at the confocal microscopy unit of the Research Infrastructure Centre at the University of Otago, Dunedin, New Zealand. This work was supported by grants from the University of Otago Research Committee and the Health Research Council of New Zealand (2/296) awarded to K. Ireton and M. Bostina and a grant from the Canadian Institutes of Health Research (CIHR) to B. B. Finlay (FDN-159935). P. Dahanayake is a recipient of a doctoral scholarship from the University of Otago.

## MATERIALS AND METHODS

### Bacterial strains and mammalian cell lines

Two variants of the wild-type (WT) enteropathogenic *E. coli* strain O127:H6 E2348/69 were used in this study (54, 55). These variants are resistant to either streptomycin (Sm^R^) or nalidixic acid (Nal^R^) (7, 31). Mutant strains of EPEC E2348/69 containing in-frame deletions of the genes *eae*, *espF*, *espG*, *espH*, *espL*, *escN*, *map*, *nleA*, *nleB1*, *nleB2*, *nleC*, *nleD*, *nleE*, *nleH1*, *nleJ*, and *tir* were previously described (7, 30, 31, 47, 53, 56–59). A strain with an in-frame deletion in *nleG* was generated using plasmid pRE112 as previously detailed (59, 60). Strain JPN15, which lacks a BFP, and isogenic strains deleted for *espH* or *tir* were described in previous work (33, 40, 61). JPN15 Δ*tir* strains harboring pACYC184 plasmids expressing wild-type *tir* or *tir* with a Y474F mutation were described previously (62). For complementation of strain JPN15 Δ*espH*, primers espH-F (5’-ATGTCGTCATCATTATCAGG-3’) and espH-R (5’-AACTGTCACACCTGATAAAG-3’) were used to PCR amplify *espH* from wild-type JPN15. The PCR product was purified using a NucleoSpin Gel and PCR clean-up kit (Machery-Nagel) and amplified using primers espHc-F (5’- CCCGTCCTGTGGATCATGTCGTCATCATTATCAGGT-3’) and espHc-R (5’- AAGGGCATCGGTCGATTAAACTGTCACACCTGATAAAGA-3’). An In-Fusion cloning kit (Takara) was used to clone the product into the BamHI/SalI sites of plasmid pACYC184 (62). The resulting pACYC184-*espH* plasmid was introduced into strain JPN15 Δ*espH* by electroporation and selection for chloramphenicol resistance.

The human cell line HeLa (ATTC CCL-2) was cultured in Dulbecco’s Modified Eagle Medium (DMEM) with 4.5 g of glucose per liter and 2 mM glutamine, supplemented with 10% fetal bovine serum (FBS) and penicillin/streptomycin.

### Antibodies

Rabbit antibodies used were anti-Sec3 (Sigma-Aldrich; HPA037706) and anti- VAMP3 (Cell Signaling Technology; 13640). Mouse monoclonal antibodies used were anti-Arp3 (Abcam; FMS338), anti-*E. coli* lipopolysaccharide (Abcam; Ab35654), anti-Exo70 (Kerafast; ED2001), anti-Exo84 (Santa Cruz Biotechnology; sc-515532), anti-EGFP clones 7.1 and 13.1 (Sigma-Aldrich; 11814460001), anti-Sec6 (Santa Cruz Biotechnology; sc-374054), anti-Sec8 (Becton Dickenson; 610658), anti- Sec10 (Abcam; ab241472), anti-Syntaxin 4 (Abcam; ab77037), and anti-tubulin (T5168; Sigma-Aldrich). Horseradish peroxidase (HRPO)-conjugated secondary antibodies were purchased from Jackson Immunoresearch. Secondary antibodies or phalloidin coupled to Alexa Fluor 488, Alexa Fluor 555, or Alexa Fluor 647 were obtained from Thermo Fisher Scientific.

*siRNAs* The sequences of short interfering RNAs (siRNAs) used were 5′-GGAAUUGAGUGGUGGUAGAtt 3′ (Arp3), 5’-CAGACAACAUCAAGAAUGAtt- 3’ (Exo70 #5), 5’-GAUUGCAUGGGCCCUUCGAtt-3’(Sec3 #4), 5’-CAGUUGUCAGGAUCAUUGAtt-3’ (Sec6 #2), 5’- CUCAUUAAGGGCUUGGCGAtt-3’ (Sec8 #2), 5’- GUUUAAUUUAGGUACUGAUtt-3’(Sec10 #1), 5’- GAGUUAACGUGGACAAGGUtt-3’ (VAMP3 #1),and 5’-GCAAUUCAAUGCAGUCCGAtt-3’(Stx4 #3). These siRNAs were obtained from Sigma-Aldrich. Non-targeting control siRNAs were purchased from Dharmacon (catalog no. D-001210-01) or Sigma-Aldrich (catalog no. SIC-001). These siRNAs have two or more mismatches with all sequences in the human genome, indicating that they should not target host mRNAs.

### Mammalian expression plasmids

Mammalian expression vectors used were pEGFP-actin (63), pEGFP-Exo70 (Addgene number 53761; gift of Channing Der), pEGFPC1 (Clontech), and pVAMP3-EGFP (14).

### Transfection

HeLa cells grown on 22- by 22-mm coverslips in 6-well plates were transfected with siRNAs or plasmid DNA using Lipofectamine 2000 (Thermo Fisher Scientific; 11668019), as described previously (28, 64).

### Western blotting

HeLa cells were solubilized in radioimmunoprecipitation assay (RIPA) buffer (1% Triton X-100, 0.25% sodium deoxycholate, 0.05% SDS, 50 mM Tris-HCl [pH 7.5], 2 mM EDTA, 150 mM NaCl, 1mM phenylmethylsulfonyl fluoride, and 10 mg/liter each of aprotinin and leupeptin) approximately 48 h post-transfection with siRNAs. Protein concentrations of lysates were determined using a bicinchoninic acid (BCA) assay kit (Pierce), and equal protein amounts of each sample were migrated on 8% or 12% SDS-polyacrylamide gels. Transfer of proteins to PVDF membranes, incubation with primary antibodies or secondary antibodies coupled to horseradish peroxidase, and detection using Clarity Max Western enhanced chemiluminescence (ECL) substrate (Bio-Rad Laboratories) were performed as described previously (65). Chemiluminescence was detected using an Odyssey imaging system (Li-Cor Biosciences). Bands in Western blot images were quantified using ImageJ software as described (66).

### Infections

EPEC strains were prepared for infection by growing overnight cultures in Luria Bertani (LB) broth at 37°C without shaking. Twenty microliters of BFP-positive EPEC strains or 40 microliters of BFP-negative JPN15 strains were then added per well. In the case of BFP-positive strains, cells were incubated for 3 h at 37°C in 5% CO_2_ in the absence of antibiotic, followed by washing in DMEM, and incubation for another 3 h in DMEM containing 100 µg/mL gentamicin. For JPN15 strains, cells were incubated for 6 h in DMEM without antibiotic. Cells were washed in DMEM at 3, 4, and 5 h post-infection.

### Imaging of exocytosis

HeLa cells were transiently transfected with plasmids expressing VAMP3-EGFP or actin-EGFP for ∼ 24 h and then infected with EPEC strains as detailed above. After infection, cells were washed twice in cold phosphate buffered saline (PBS) and subjected to exofacial labeling of VAMP3-EGFP using mouse anti-GFP antibodies, as previously described (21, 22, 28). Cells were then fixed in PBS containing 3% paraformaldehyde (PFA), quenched in PBS with 50 mM ammonium chloride, and labeled with anti-mouse-Alexa Fluor 647 antibodies to detect exofacial VAMP3- EGFP. Samples were permeabilized in PBS containing 0.4% Triton X-100 and incubated with phalloidin coupled to Alexa Fluor 555 to label F-actin and the DNA stain DAPI to detect bacteria. Samples were mounted in Molwiol® supplemented with DABCO (1,4-diazabicyclo[2.2.2]octane).

Imaging was performed using Zeiss LSM900 or Olympus FV1200 laser scanning confocal microscopes equipped with 63x or 60x 1.4 NA oil immersion objectives. Laser lines of 405, 488, 561, and 640 nm, were used for excitation and GaAsP photocathodes or photomultiplier tubes were used for detection. Images from serial sections spaced 0.75 µm apart were used to ensure that all pedestals were detected. Pedestals were identified as actin-rich structures projecting from the apical surface of cells. Details on the numbers of pedestals imaged are provided in the figure legends. The threshold function of ImageJ2 (version 2.16.0/1.54p) software was used to determine fold enrichment (FE) values for exofacial VAMP3-EGFP for each pedestal, as described in previous work with protrusions made by *Listeria* or *Shigella* (21, 22). FE values were determined as the mean pixel intensity of VAMP3-EGFP in a pedestal divided by the mean pixel intensity throughout the entire host cell.

### Analysis of recruitment of Exo70

For studies with EGFP-Exo70, HeLa cells were transfected with plasmids expressing human Exo70 containing an amino-terminal EGFP tag. As a control, cells were transfected with a plasmid expressing EGFP alone. About 24 h post-transfection, HeLa cells were infected with WT, Δ*tir,* or Δ*espH* strains of JPN15 for 6 h (Figs. 4 and 9B), followed by fixation in PBS with 3% PFA, quenching in PBS with 50 mM ammonium chloride, and permeabilization in PBS with 0.4% Triton X-100. Samples were incubated with DAPI to detect DNA and phalloidin coupled to Alexa Fluor 555 to label F-actin. Experiments detecting endogenous Exo70 (Figure S4) were performed similarly, except that HeLa cells not transfected, and Exo70 was labeled using mouse anti-Exo70 antibodies. Samples were imaged by confocal microscopy and the degree of recruitment of EGFP-Exo70 or EGFP was quantified as FE values, similarly to as described for FE quantification of exofacial VAMP3-EGFP.

### Analysis of distribution of exocytosis or Exo70 around pedestals

For assessment of exocytosis distribution, HeLa cells transiently expressing VAMP3- EGFP were infected with WT or Δ*espH* JPN15 strains for 6 h, subjected to exofacial labeling, fixed, permeabilized and labeled with phalloidin-AlexaFluor555 to detect F-actin and with DAPI to stain DNA (Figure 10A and supplemental movies 1 and 2). Analysis of EGFP-Exo70 localization (Figure S6A and supplemental movies 3 and 4) was performed similarly to the exocytosis studies, except that samples of HeLa cells transiently expressing EGFP-Exo70 were used. A Zeiss LSM900 confocal microscope operating in the LSM Plus mode was used to image approximately 60 optical sections placed 0.23 µm apart. ZEISS arivis Pro software (version 4.2.1) was used to assemble optical sections and generate movies.

### Quantification of pedestal formation

For experiments involving depletion of exocyst or SNARE proteins by RNAi (Figures 3, 6, and S2), HeLa cells were either mock transfected in the absence of siRNA, transfected with 100 nM of control siRNA, or transfected with the same concentration of siRNAs targeting Sec3, Sec6, Sec8, Sec10, Exo70, VAMP3, or Stx4. Experiments to co-deplete Arp3 with Sec3, Exo70, or VAMP3 (Figure 7), were performed similarly, except that cells were co-transfected with 50 nM of each siRNA. About 48 h after transfection, cells were infected with wild-type JPN15 for 6 h, washed in PBS, fixed in PBS containing 3% PFA, quenched in PBS with 50 mM ammonium chloride, and permeabilized in PBS with 0.4% Triton X-100. Infected HeLa cells were incubated with phalloidin conjugated to Alexa Fluor 555 to label F-actin in pedestals and DAPI to detect DNA. To assess pedestal formation efficiencies, confocal microscopy images were obtained at 0.75 µm intervals. Bacteria associated with host cells were identified by both DAPI labeling and differential interference contrast microscopy (DIC). Efficiencies of pedestal formation were initially determined as the percentage of cell-associated bacteria that had pedestals or the percentage of HeLa cells with more than 5 pedestals. In Figures 3B and 7E, the resulting percentages were then converted to relative pedestal formation values by normalization to the percentage value for the no siRNA condition. In parallel with preparing samples for confocal microscopy analysis of pedestal formation, HeLa cells subjected to the same siRNA transfection conditions were solubilized in RIPA buffer to perform Western blots to confirm depletion of target proteins (Figures S2 and 7B,C,D). Experiments comparing pedestal formation efficiencies of WT and Δ*espH* strains of JPN15 (Figures 9C and S5C) were performed similarly to as described above, except that HeLa cells were not subjected to siRNA transfection.

### Quantification of pedestal, exocytosis, or Exo70 volumes

For studies comparing WT and Δ*espH* strains of JPN15 (Figures 10B and S6B), HeLa cells transiently expressing VAMP3-EGFP or EGFP-Exo70 were infected with bacteria for 6 h. Cells with VAMP3-EGFP were subjected to exofacial labeling to detect exocytosis. Samples were then fixed, permeabilized, and labeled with phalloidin-AlexaFuor555 to detect F-actin in pedestals. Experiments to assess the effects of RNAi-mediated depletion of Sec3 or Stx4 (Figure 11) were performed similarly, except that HeLa cells were transfected with control siRNA or siRNAs targeting Sec3 or Stx4 about 24 h prior before transfection with plasmids expressing VAMP3-EGFP, and ∼ 48 h prior to infection with WT JPN15. A Zeiss LSM900 microscope equipped with an Airyscan 2 super-resolution module was used to obtain images of optical sections spaced 100 nm apart. Images were further processed by deconvolution and Zeiss arivis Pro software to perform AI-assisted segmentation and 3D reconstruction. Volumes of pedestals were quantified using reconstructed images of F-actin. Volumes of exocytosis or Exo70 associated with pedestals were also determined.

### Analysis of pedestal lengths and areas

HeLa cells were infected with a wild-type Sm^R^ BFP-positive strain of EPEC O127:H6 E2348/69 or an isogenic mutant strain deleted for *espH* for 6 h. Cells were then fixed in 2.5% glutaraldehyde in 0.176 M cacodylate buffer, subjected to high pressure freezing using a Leica EM Pact2 device, embedded in epoxy resin, and imaged using a Zeiss Sigma 300 serial scanning electron microscope equipped with a Gatan 3View2XP serial block face system. Lengths and areas of pedestals were measured using Microscopy Image Browser (MIB) software and areas were measured using ImageJ2 software.

### Statistical analysis

Statistical analysis was performed using Prism (version 8.4.3; GraphPad Software). In comparisons of data from three or more conditions, analysis of variance (ANOVA) was used. The Tukey-Kramer test was used as a posttest. For comparisons of two conditions, an unpaired t-test was used. A P-value of 0.05 or lower was considered significant.

